# Sexual Dimorphism of Plasma and Tissue Proteomes in Human Calcific Aortic Valve Stenosis Pathogenesis

**DOI:** 10.1101/2024.12.09.627613

**Authors:** Cassandra L. Clift, Mark C. Blaser, Francesca Bartoli-Leonard, Florian Schlotter, Hideyuki Higashi, Shiori Kuraoka, Taku Kasai, Mandy E. Turner, Tan Pham, Katelyn A. Perez, Simon C. Robson, Simon C. Body, Jochen D. Muehlschlegel, Masanori Aikawa, Sasha A. Singh, Elena Aikawa

## Abstract

**BACKGROUND:** Calcific aortic valve stenosis (CAVS) is a global clinical burden, impacting around 2% of the population over 65 years of age. No pharmacotherapeutics exist, with surgical repair and transcatheter valve replacement being the only intervention. Females are underrepresented in studies of CAVS, leading to delay in timely intervention and increased mortality. Histopathology demonstrates female CAVS presents with decreased valvular calcification but increased fibrosis and severity of symptoms. We hypothesize that the underlying molecular mechanisms contributing to disease progression and fibrocalcific burden in AS differs between male and female patients. Our goal for this study is to use previously acquired proteomic datasets of a clinically-defined human AS cohort to examine sex disparities and underlying sex-specific disease signatures.

**METHODS and RESULTS:** Age-matched human AS tissue samples (n=4 females, n=14 males) were each segmented into non-diseased, fibrotic, and calcified disease stages and analyzed using LC-MS/MS proteomics and quantitative histopathology. CAVD plasma samples (n=20 females, n=30 males) were analyzed for circulating sex-specific biomarkers via LC-MS/MS. Unbiased principal component analysis shows sex- and stage-specific proteome clustering. AS pathogenesis drove sex-specific disparities in the valvular proteome: 338/1503 total proteins were differentially-enriched by sex across disease stages. Compared to sex-specific non-diseased controls, female fibrotic tissue resulted in 2.75-fold greater number of differentially-enriched proteins than did male fibrotic tissue (female: 42, male: 16; p<0.05 threshold). In contrast, female calcific tissue identified 2.473-fold less differentially-enriched proteins than male calcific tissue (female: 157, male 356; q<0.05 threshold). By Functional Enrichment Analysis revealed specific proteins responsible for the exacerbated valvular fibrosis signature in females, implicated adenosine phosphate metabolism as a potential male-specific driver of AS, and further reinforce the shared contribution of aberrant lipid and cholesterol activity to AS progression in both sexes.

**CONCLUSIONS:** We reveal a sexually-dimorphic AS proteome, including the novel overabundance of ECM remodeling pathways in female calcified aortic valve tissues. This analysis allows for identification of potential sex-specific protein drug targets implicated in AS pathobiology.

## Manuscript

Calcific aortic valve disease resulting in aortic stenosis (AS) impacts 2% of those over age 65, causing heart failure as the aortic valve (AV) becomes fibrotic, calcified, and obstructs cardiac function. No pharmacotherapies exist for AS, with surgical/transcatheter valve replacement being the only treatment option. There are glaring sex disparities in the clinical presentation and treatment of AS. Females have decreased likelihood to receive timely intervention, worse symptoms, poorer NYHA scoring, and increased mortality rates during AV replacement^1^. Although seminal histopathological evaluations found that females have elevated valvular fibrosis while incurring similar hemodynamic deficits to males^2^, few molecular studies of underlying sexual dimorphism in AS exist. To discover sex-specific biomarkers and pharmacotherapeutic targets, it is critical to identify sexually dimorphic molecular drivers of fibrocalcific disease progression.

Here, we report the first proteomic fingerprint of human AS sex disparities in patients undergoing surgical AV replacement for moderate-to-severe hemodynamic dysfunction. We reveal sexual dimorphism in the circulating plasma proteome of AS patients using novel mass spectrometric plasma proteomics, and perform *post-hoc* sex-stratified analysis of AS histopathological and tissue proteomics datasets^3^. In tissue-based analyses, stenotic AV leaflets were segmented into non-diseased, fibrotic, and calcified disease stages. Extensive validation (histopathology, near-infrared molecular imaging, electron microscopy, proteomics/transcriptomics) has previously shown that non-diseased regions are comparable to healthy AVs, while disease burden continuously grows in fibrotic and calcified stages (**Figure A**)^3,4^. Together, our studies uncover molecular pathways underpinning differences in fibrocalcific AV disease progression between males and females.

**Figure 1:**
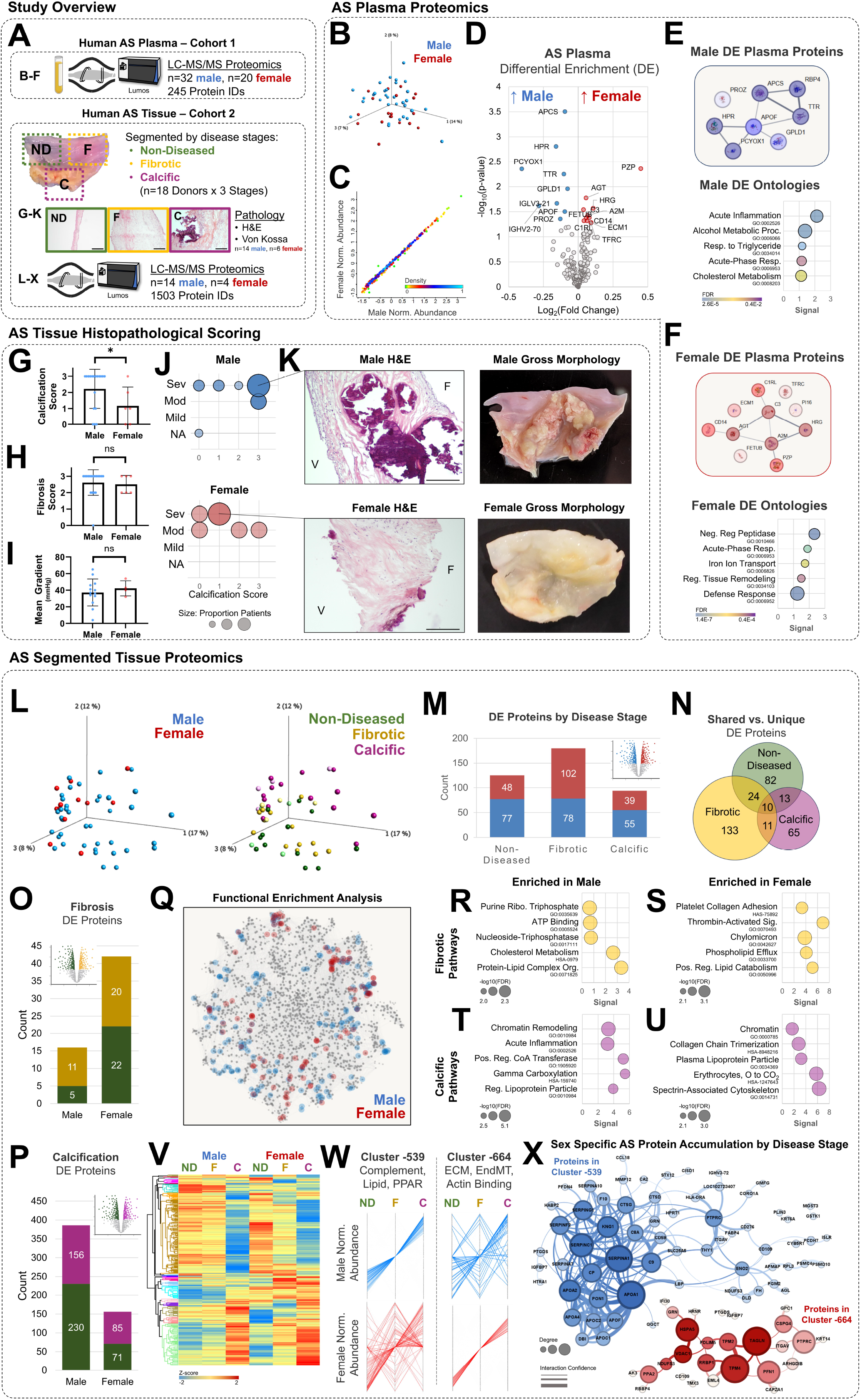
Sex-specific pathological signatures of human aortic stenosis (AS). **A)** Schematic summary of study design and acquisition strategy. Mass spectrometry based plasma and tissue proteomics were performed on two separate cohorts of patients undergoing aortic valve replacement for moderate-to-severe aortic valve stenosis. LC-MS/MS analysis was generated via data dependent acquisition on a Fusion Lumos mass spectrometer equipped with an EASY nLC 1100 system (Thermo Fisher Scientific). CID tandem mass spectra were searched using Proteome Discoverer version 2.5. **B-F: Plasma proteomics of male and female AS patients. B)** Principal component analysis (PCA) of plasma proteome in male (n=32) versus female (n=20) AS plasma. **C)** Log2 protein abundance scatterplot with density color-scale **D)** Volcano plot of differentially enriched (DE) plasma proteins in males and females; Students t*-test with p<0.05. **E-F)** String-DB protein-protein interaction (PPI) networks of differentially enriched plasma proteins in **(E)** male and **(F)** female cohorts, with corresponding enriched Gene Ontology Biological Processes (GOBP). Female PPI enrichment p=1.93e-05. Male PPI enrichment p=1.13e-13 (STRING-DB). ‘Signal’ is defined as the weighted harmonic mean of log_10_(observed/expected) and –log(FDR), and is calculated with one degree of network freedom with the second shell containing no more than 5 interactions. Bubble size is count of proteins annotated with the respective term. **G-K: Histopathological scoring of sex disparities in aortic valve fibrosis severity as a function of calcification score and sex. G)** Quantification of male (n=14) vs. female (n=6) aortic valve calcification score; *p=0.02. **H)** Quantification of male vs. female aortic valve fibrosis score; ns p>0.05. **I)** Male vs. female echocardiographic mean aortic valve gradient; ns p>0.05. Mean±SEM. **J)** Bubble plots of categorical male (top) and female (bottom) calcification score versus fibrosis severity. Highly-fibrotic female aortic valves have less calcification than their male counterparts. **K)** Representative hematoxylin and eosin (H&E) histopathological staining of longitudinal aortic valve sections from predominant patient groups (Males: calcification = 3, fibrosis = severe; Females: calcification = 1, fibrosis = severe). Representative images of male and female gross aortic valve morphology. Quantitative blinded pathologist-graded fibrosis scores (0-3, none [NA]/mild/moderate/severe) via H&E, and calcification scores (0-3, none [NA]/mild/moderate/severe) via Von Kossa (not shown). Bar = 200μm. **L-X: Disease stage segment AS tissue proteomics of male and female tissues. L)** PCAs of male and female (left) disease stage-specific (right) tissue proteomics show reduced stage-to-stage proteome variance in females. **M-N: Male vs. female differentially-enriched proteins, within each disease stage. M)** Quantification of male vs. female differential protein enrichment (DE, p≤0.05) per disease stage reveals that the highest number of DE proteins between sexes occurs within fibrotic regions of stenotic aortic valves. **N)** Venn diagram of male vs. female DE proteins per disease stage shows few sex-altered proteins are shared between stages, suggesting male and females have distinct disease progression. **O-X: Non-diseased vs. fibrotic/calcified stage differentially-enriched proteins, within each sex. O)** Quantification of non-diseased vs. fibrotic stage DE proteins in males or females (p≤0.05). **P)** Quantification of non-diseased vs. calcified stage DE proteins in males or females (p≤0.05). **Q)** PPI network of the overall AS tissue proteome (1503 total protein IDs), with highlighted log fold change ranked Functional Enrichment Analysis (FEA) of male and female between-disease stage DE proteins that were found in **O-P**. FEA incorporates Gene Ontology (GO) Biological Process, GO Molecular Function, GO Cellular Component, KEGG, and Reactome. **R-U:** Highlighted significantly enriched (FDR≤0.05) FEA ontologies and pathways in **(R)** male fibrosis, **(S)** female fibrosis, **(T)** male calcification, and **(U)** female calcification. **V)** Hierarchically clustered heat map of between-disease stage DE proteins identified in **O-P. W)** Profile plots of proteins within selected clusters from **V** with progressive protein accumulation along disease stages in only males (Cluster −539, left) and only females (Cluster −664, right). **X)** PPI networks of proteins comprising the clusters profiled in **W**. Cluster −539 PPI enrichment p<1.0E-16. Cluster −664 PPI enrichment p=1.52E-6.

We first performed plasma proteomics (n=32 males, 20 females) to screen for sex-dimorphic circulating biomarkers of AS (**Figures B-F**). Despite high male/female correlation of the plasma proteome (**Figures B-C**), 20 proteins were differentially-enriched between sexes (**Figure D**). Protein-protein interaction (PPI) networks and pathway enrichment found that male-enriched plasma proteins were associated with acute inflammation (APCS, SAA2-4, APOA2, HPR; FDR=2.6e-5) and cholesterol metabolism (APOL1/A4/A2/F; FDR=3.9e-3) (**Figure E**), while metalloendopeptidase activity (HRG, C3, PZP, FETUB, A2M, AGT, ECM1, PI16; FDR=1.38e-7) and tissue remodeling (HRG, AGT, TFRC, TF; FDR=3.9e-4) were female-enriched (**Figure F**). Increased angiotensinogen (AGT) in female AS plasma is notable, as angiotensin receptor blockers were recently found to be negatively associated with valvular fibrosis only in female AS, through an as-yet-undetermined mechanism-of-action^5^.

In AV tissues, male AS leaflets (n=14) had significantly higher calcific burden than females (n=6) (**Figure G**, p=0.0212), while fibrosis severity (**Figure H**) and mean AV gradient (**Figure I**) did not differ between sexes – consistent with prior reports^2^ (**Figure H**). Categorical analysis found that highly-fibrotic female AVs have less calcification than their male counterparts (**Figure J-K**). Amongst 22 clinical characteristics, only incidence of hypertension (p=0.004) and beta-blocker usage (p=0.0233) differed between males (both elevated) and females in our tissue cohort. Male vs. female disease stage-specific tissue proteomics then identified proteins and pathways contributing to this sexually dimorphic histopathological presentation of AS (**Figure L-X**). Principal component analysis revealed sex- and stage-specific tissue proteome clustering (**Figure L**). AS pathogenesis drove sex-specific disparities in the valvular proteome, with fibrotic segments containing the largest number of differentially-enriched proteins (180/399 DE proteins) (**Figure M**). Differentially-enriched proteins were predominantly unique across disease stages, suggesting sex-dimorphic disease progression (**Figure N**).

Within each sex, proteomic comparisons across diseases stages (**Fibrotic:** Figure O; Calcified: **Figure P**) exposed sex- and stage-specific propensities in differential enrichment: females had 2.75-fold more altered proteins in fibrosis, while males had 2.47-fold more in calcification. Functional Enrichment Analysis identified ubiquitous cholesterol metabolism,phospholipid efflux, and lipoprotein particles in both sexes during AS pathogenesis (**Figures Q-U**). In fibrosis, males are uniquely enriched in nucleoside triphosphate pathways – suggesting sex-specificity of established roles for ATP catabolism in AS (**Figure R**), along with inflammatory pathways in calcification (**Figure T**). Unique enrichment of platelet-collagen adhesion (**Figure S**) and collagen chain trimerization (**Figure U**) uncovered a specific basis for the highly-fibrotic nature of stenotic female AVs. Profiles of proteins with pathogenic accumulation (**Figure V-W**) in only males (Cluster-539) or females (Cluster-664) and PPIs (**Figure X**) showed male AS across progressive disease stages is defined by complement activation, lipid accumulation, and PPAR signaling (SERPINA1/C1/F2/A7/G1, APOA1/A2/A4/C2/C1/F; p<1.0E-16), while female AS pathogenesis is distinguished by extracellular matrix matricrine signaling, endothelial-to-mesenchymal transition, and actin binding (ITGAV, IGFBP7, ECM2, SPARCL1; p<1.52E-6).

In conclusion, this is the first proteomics study of sexual dimorphism in human AS tissues, and the first report of mass spectrometry-based plasma proteomics in human AS of any kind. Previous studies of sex differences in AS employed histopathology^2^ which cannot pinpoint a molecular-level disease fingerprint, or transcriptomics^4^ which do not correlate well with tissue-level protein signatures in the matrix-dense and cell-sparse AV. Here, we reveal a sexually-dimorphic AS proteome and pinpoint putative circulating sex-specific AS biomarkers. Importantly, we identify specific fibrotic matrix constituents and metabolism pathways unique to female AS pathogenesis. These findings potentiate future studies aimed at identifying male- and female-tailored therapeutic targets and treatment strategies in this as-yet-intractable disease^5^.

## Acknowledgements

We thank Johana Barrientos and Daniel Deppe for assistance with histology.

## References

1. DesJardin JT, Chikwe J, Hahn RT, Hung JW, Delling FN. Sex Differences and Similarities in Valvular Heart Disease. Circulation Research 2022;130:455–473. doi:10.1161/CIRCRESAHA.121.319914.

2. Simard L, Côté N, Dagenais F, Mathieu P, Couture C, Trahan S, et al. Sex-related discordance between aortic valve calcification and hemodynamic severity of aortic stenosis. Circulation Research 2017;120:681–691. doi:10.1161/CIRCRESAHA.116.309306.

3. Blaser MC, Buffolo F, Halu A, Turner ME, Schlotter F, Higashi H, et al. Multiomics of Tissue Extracellular Vesicles Identifies Unique Modulators of Atherosclerosis and Calcific Aortic Valve Stenosis. Circulation 2023;148:661–678. doi:10.1161/CIRCULATIONAHA.122.063402.

4. Sarajlic P, Plunde O, Franco-Cereceda A, Bäck M. Artificial Intelligence Models Reveal Sex-Specific Gene Expression in Aortic Valve Calcification. JACC Basic Transl Sci 2021;6:403– 412. doi:10.1016/j.jacbts.2021.02.005.

5. Carter-Storch R, Nezet EL, Ali M, Powers A, Haujir A, Demers K, et al. Angiotensin II Receptor Blockers Are Associated With Reduced Valvular Fibrosis in Women With Aortic Stenosis. Canadian Journal of Cardiology 2024;40:1690–1699. doi:10.1016/j.cjca.2024.03.009.

